# Spatially repeatable components from ultrafast ultrasound are associated with motor unit activity in human isometric contractions

**DOI:** 10.1101/2023.04.17.537211

**Authors:** Robin Rohlén, Marco Carbonaro, Giacinto L. Cerone, Kristen M. Meiburger, Alberto Botter, Christer Grönlund

## Abstract

**Objective:** Ultrafast ultrasound imaging has been used to measure intramuscular mechanical dynamics associated with single motor unit (MU) activations. Detecting MU activity from ultrasound sequences requires decomposing a displacement velocity field into components consisting of spatial maps and temporal displacement signals. These components can be associated with putative MU activity or spurious movements (noise). The differentiation between putative MUs and noise has been accomplished by comparing the temporal displacement signals with MU firings obtained from needle EMG. Here, we examined whether the repeatability of the spatial maps over brief time intervals can serve as a criterion for distinguishing putative MUs from noise in low-force isometric contractions.

**Approach:** In five healthy subjects, ultrafast ultrasound images and high-density surface EMG (HDsEMG) were recorded simultaneously from biceps brachii. MUs identified through HDsEMG decomposition were used as a reference to assess the outcomes of the ultrasound-based decomposition. For each contraction, displacement velocity sequences from the same eight-second ultrasound recording were separated into consecutive two-second epochs and decomposed. The Jaccard Similarity Coefficient (JSC) was employed to evaluate the repeatability of components’ spatial maps across epochs. Finally, the association between the ultrasound components and the MUs decomposed from HDsEMG was assessed.

**Main results:** All the MU-matched components had JSC > 0.38, indicating they were repeatable and accounted for about one-third of the HDsEMG-detected MUs (1.8 ± 1.6 matches over 4.9 ± 1.8 MUs). The repeatable components (with JSC over the empirical threshold of 0.38) represented 14% of the total components (6.5 ± 3.3 components). These findings align with our hypothesis that intra-sequence repeatability can differentiate putative MUs from spurious components and can be used for data reduction.

**Significance:** The results of our study provide the foundation for developing stand-alone methods to identify MU in ultrafast ultrasound sequences and represent a step forward towards real-time imaging of active MU territories. These methods are relevant for studying muscle neuromechanics and designing novel neural interfaces.

## Introduction

Recently, neuromuscular imaging based on ultrafast ultrasound (UUS) has evolved considerably, opening new fronts in studying muscle contraction at the single motor unit (MU) level [1–9]. High-resolution imaging of active muscle tissue can provide spatiotemporal mechanics of individual MU fibres, complementing the information accessible with standard electrophysiological techniques for assessing single MU properties, i.e., invasive needle electromyography (nEMG) [10–12] and non-invasive surface EMG (sEMG) [13,14]. The added information on spatial and temporal mechanics can foster basic studies on muscle neuromechanics and force generation mechanisms [15], along with providing biomarkers for myopathic disorders [16–18], and innovative neural interfaces relevant, e.g., in rehabilitation and prosthetic control [19–21].

The methodology of identifying single MU activity in UUS recordings during isometric *voluntary* contractions was recently proposed based on a two-step approach [3]. First, the subtle intramuscular displacement velocities were estimated [22], and then these displacement velocities were decomposed into multiple components. Each component comprises a *spatial* map (location of the component, related to MU territory) and a *temporal* signal (time course of its displacement velocity, related to MU spike train). To separate spurious components (noise) from those associated with single MU activation, a procedure based on *temporal* signal characteristics was adopted and later validated against single MU identification based on needle EMG [4]. It was found that a large proportion of the components’ temporal twitch-by-twitch signals could not be matched with MU firings [4,6]. Two factors may contribute to this relatively low agreement between the two measures. The first is the heterogenic composition of linear and non-linear elastic tissue constituents, causing a non-linear combination of MU twitches. The second one concerns MU firing variability. Indeed, although the MU pool should be stable during these contractions, the firing rate of MUs varies, which has been shown to influence the temporal twitch parameters, i.e., alter the temporal signal (sequence of twitches) [15].

In contrast to the temporal firing characteristics, the location of MU fibres within the muscle cross-section should represent an invariant feature during constant force and isometric contractions. It follows that components with a stable spatial map throughout the contraction are more likely to be associated with actual MU activations. Hence, we hypothesise that the spatial repeatability of a component across short epochs (intra-sequence repeatability) is a feature associated with MU activity and may be used as a criterion for data reduction of the initial decomposed components. In this study, we aimed to identify intra-sequence spatially repeatable components and examine whether repeatability can be used to separate MUs from noise in stable low-force isometric contractions. For this purpose, we decomposed displacement velocity images in consecutive two-second epochs from eight-second UUS recordings. We quantified the repeatability of the components’ spatial map across epochs and examined whether the repeatable components were associated with actual MU activity. To this end, we used a set of reference MUs identified with an independent and validated decomposition method (HDsEMG decomposition [23]), applied to experimental signals detected simultaneously with the ultrasound images. Finally, we determined whether the analysis based on two-second intervals (required to assess the repeatability) affects the number of MU-matched components compared with the decomposition of the recordings’ full length (eight seconds).

## Methods

### Experimental protocol

Five subjects (31 ± 6 years, three males, and two females) performed three low-level isometric constant-force elbow flexions (from 2% to 10% of the maximum voluntary contraction). The details of the experimental protocol are reported in Carbonaro et al. [6]. Briefly, for each contraction, eight-second-long UUS recordings (Verasonics Vantage 128, Verasonics Inc., Kirkland, WA) were recorded simultaneously [24] with HDsEMG (MEACS, LISiN, Politecnico di Torino, Turin, Italy [25]). A grid of 64 surface-EMG electrodes transparent to ultrasounds (8×8, 10 mm inter-electrode distance [26]) was placed on the muscle belly with the ultrasound transducer (L11-5v, 7.81 MHz centre frequency, 31.25 MHz sampling rate, and 2500 Hz frame rate) positioned between the fourth and the fifth row of electrodes; i.e. transversally with respect to the muscle fibres’ direction (Fig. 1A). The study was conducted following the Declaration of Helsinki and approved by the Regional Ethics Committee. Informed consent was obtained from all subjects.

**Figure 1.**
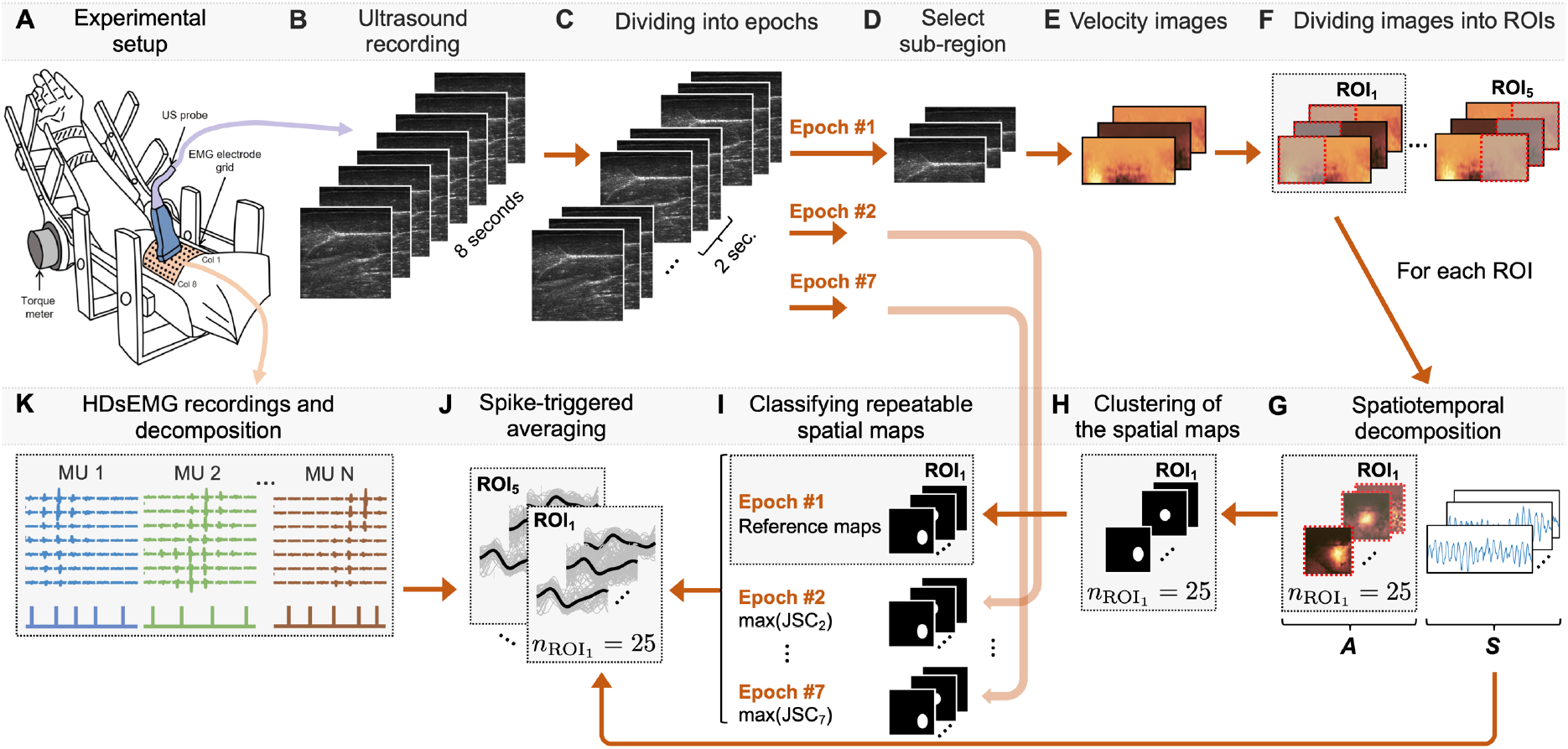
Illustration of the ultrasound data processing and identification of repeatable spatial maps. **A**. Experimental setup with simultaneous ultrafast ultrasound (UUS) and high-density surface electromyography (HDsEMG) recordings (adapted from Carbonaro et al. (2022) [6]). **B**. Eight-second recordings using UUS (40×40 mm, 2500 Hz) plane wave imaging. **C**. The recordings were divided into seven partially overlapping epochs of two seconds each. **D**. A sub-region was selected within the HDsEMG detection volume (20×40 mm). **E**. Tissue velocity images were estimated. **F**. The velocity images were divided into five region-of-interests (ROIs), i.e., 20×20 mm each. **G**. Each ROI was decomposed into 25 components, i.e., 25 temporal signals (‘S’) and 25 spatial maps (‘A’). **H**. The spatial maps were clustered and processed to generate a binary map, with zeros being the background and ones being the largest intensity of the territory. **I**. The binary maps were used for calculating the Jaccard Similarity Coefficient (JSC) for each component in the epoch (second to the seventh) with the first epoch as a reference. The maximal JSC was retained for each epoch, and then the mean JSC (based on the maximal JSC for all epochs) was calculated. **J**. Then, spike-triggered averaging of the components’ temporal signal was performed using the motor unit (MU) spike trains instants from the **K**. HDsEMG decomposition.

### UUS and HDsEMG data processing

The radio frequency UUS data comprised 20000 frames (2176×128 pixels, i.e., approximately 53×40 mm). After traditional delay-and-sum beamforming, each eight-second dataset was processed in two-second epochs [3,4] with one-second overlapping ([0:2] s, [1:3] s, …, [5:7] s, [6:8] s) resulting in seven sub-datasets of two seconds (Fig. 1C). Each pixel in each sub-dataset was filtered over time with a 1D median filter with the order equal to 10 ms [3,4]. The image was cropped to 20×40 mm (850×128 pixels) [6,7] (Fig. 1D). For each epoch, displacement velocity images were calculated using 2D autocorrelation velocity tracking [22,27] with 1 mm in-depth and a sliding window of 10 ms (Fig. 1E). The temporal evolution of each pixel in the velocity images was high pass filtered at 3 Hz using 3^rd^ order Butterworth filter (zero-phase) to attenuate slow movements not associated with muscle contraction [3]. Finally, the velocity images were down-sampled to 63×128 pixels, corresponding to approximately 0.3×0.3 mm per pixel.

HDsEMG signals were bandpass filtered (20-400 Hz) and decomposed into individual MU spike trains [23] (Fig. 1K). The spike trains were edited [28] and resampled at the ultrasound frame rate. MU action potential (MUAP) amplitude distributions and their centroids were calculated using the longitudinal single differential MUAP decomposed from HDsEMG [29]. Considering that the mediolateral surface covered by the HDsEMG grid is larger than that of the ultrasound transducer (Fig. 1A), all the centroids with the mediolateral coordinate outside the ultrasound field of view were truncated to the position of the first or last element of the probe (i.e., element 1 or 128).

### Spatiotemporal decomposition of displacement velocity images

As described in previous papers, the displacement velocity images were processed over five partially overlapping Region of Interest (ROIs) of 20×20 mm (5 mm increments) [4,6,8] (Fig. 1F). We used spatiotemporal independent component analysis (stICA) [30] with α = 1.0 [8] to obtain 25 spatial components (*spatial maps*) and corresponding temporal components (*temporal signals*) per ROI [4,8] (Fig. 1G). Hence, we obtained 125 spatiotemporal *ultrasound components* for each recording.

We clustered the intensities of each spatial map using the k-means algorithm with five clusters based on Euclidean distance (Fig. 1H). The cluster with the highest intensity values was assumed to be the localised spatial region (territory) of interest. Given this cluster, a binary map was generated. Objects with less than 25 connected pixels (∼1.5×1.5 mm^2^) were removed to remove noisy pixels at other regions in the image.

### Repeatability analysis: selecting similar spatial maps across epochs

A Jaccard Similarity Coefficient (JSC) criterion based on the binary maps was used to select a set of similar spatial maps across different time epochs. Specifically, the 25 spatial maps of the *first two-second epoch* for each ROI were regarded as *reference* maps (Fig. 1I). Jaccard Similarity Coefficients were calculated between each *reference* map and the 25 maps obtained from each of the remaining six epochs. For each epoch, the map with the highest JSC was retained. This procedure provided, for each *reference* map, a selection of six spatial maps maximally similar to it. The *mean spatial map* and *mean JSC* (indicating the level of repeatability of a component) were then computed using the selected maps. In total, 25 mean spatial maps were identified for each of the five ROIs (125 mean spatial maps, including all five ROIs).

### Association of selected similar components with MUs from HDsEMG

We studied the association between the ultrasound components selected in the previous paragraph and the characteristics of individual MUs identified through HDsEMG decomposition. To this end, we considered the *temporal* signal corresponding to the selected spatial maps and the firing pattern of the MUs identified from HDsEMG.

The *temporal* signals of each set of selected components were spike-triggered averaged (Fig. 1J) using the spike train of individual MUs identified from HDsEMG (Fig. 1K). This procedure was applied to all the combinations of selected ultrasound components and HDsEMG MUs, leading to a large set of *putative twitches* (Fig. 1J). Only those whose peak-to-peak amplitude exceeded a noise threshold were retained among these putative twitches. Among this subset, the pair (ultrasound component – HDsEMG MU) leading to the highest twitch amplitude was called the *MU-matched* component. The noise threshold was calculated by generating 125 temporal components of coloured noise (5-30 Hz bandwidth of white noise) and spike-triggered averaged with 100 random spike trains (mean firing rates between 8-20 Hz and standard deviation of 15% of the mean inter-pulse interval [31]). The threshold value was computed as the mean plus two standard deviations of the peak-to-peak amplitudes of all the combinations of random components and spike trains.

### Number of matched components with MUs from HDsEMG: intra and full sequence approach

We intended to assess whether the analysis on two-second intervals, required to assess the repeatability, affected the number of MU-matched components. Therefore, we compared the number of MU-matched components found with the *intra-sequence* repeatability approach with the components decomposed from the stICA applied over the *full sequence recording* [4]. In both approaches, the matching with HDsEMG MUs was performed using the same method described in the previous paragraph.

### Statistical analysis

We calculated descriptive statistics associated with the components (epochs and full sequence) and the MUs decomposed from HDsEMG. Based on the matched components with MU, we calculated the area, equivalent diameter (square root of 4xArea/π as in [3]), and depth of the centroid of the component below the skin. In addition, the distance between the mediolateral centroids of the spatial map (based on the binary map) and MUAP spatial distribution (based on the spike-triggered average on the HDsEMG signals using the MU spike trains [29]) for each matched component and MU was calculated.

We tested the pairwise difference between the number of MU-matched components between the intra-sequence repeatability and the full sequence approach using a two-sided Wilcoxon signed rank test. In addition, we tested the difference in median JSC and normalised peak-to-peak amplitude, respectively, between the MU- and non-MU-matched components using the Mann-Whitney U test. The significance level was set to 0.05.

## Results

Out of 20 recordings, 99 MUs (4.9 ± 1.8 MUs per recording) were identified by decomposing HDsEMG signals. The MUs had stable spike trains over the eight-second recordings with firing rates of 12.3 ± 2.1 Hz.

We observed various degrees of intra-sequence repeatability across the 125 ultrasound components per recording, as shown by the large variability of JSC values (Fig. S1 in Supplementary material). Fig. 2 depicts two examples of repeatable components (high mean JSC) and one non-repeatable component (low mean JSC) from one ROI of a representative subject recording.

**Figure 2.**
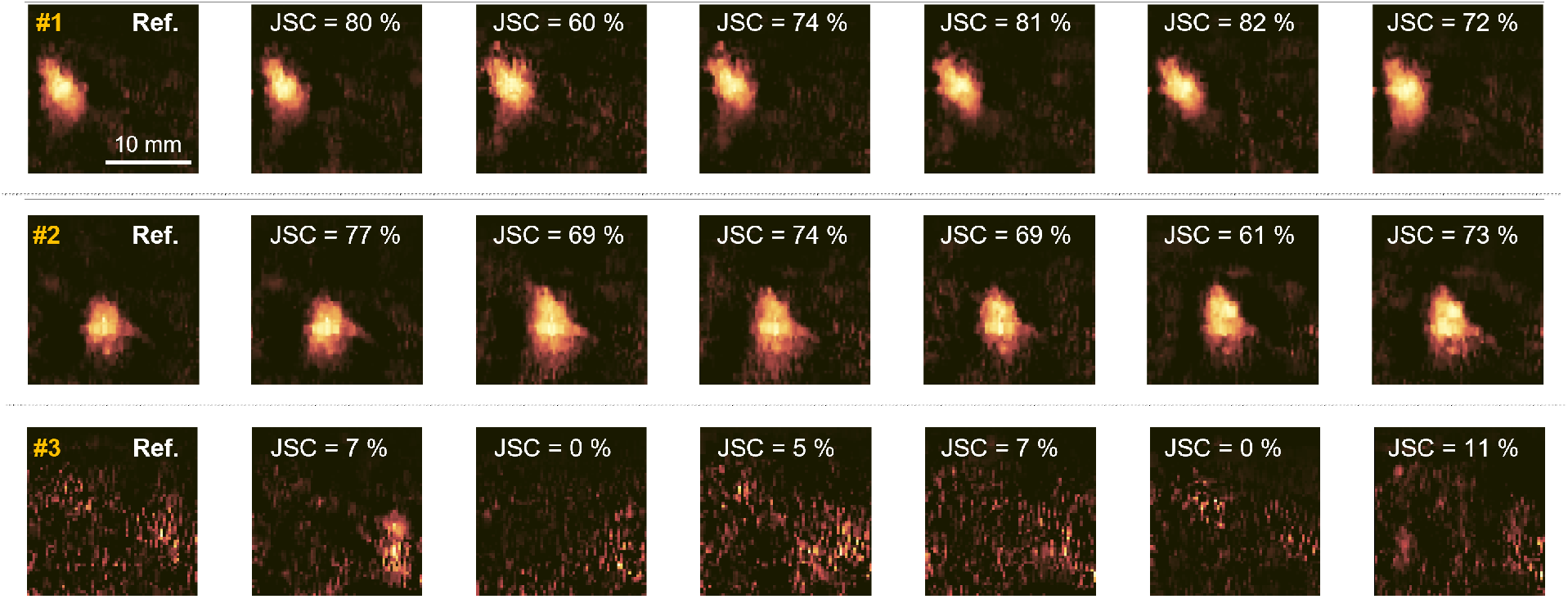
Examples of repeatable spatial maps from two repeatable components (#1 to #2) and one non-repeatable component (#3) of the same recording and region-of-interest (ROI) based on the Jaccard Similarity Coefficient (JSC). The first two-second epoch is the reference (defined as Ref).

### Association of selected similar components with MUs from HDsEMG

The scatterplot of Fig. 3 shows the relationship between JSC values and the amplitude of the (spike-triggered averaged) *putative twitches* from all subjects and trials. Each data point in Fig. 3 represents an ultrasound component and an HDsEMG MU that provided the *putative twitch* with the highest amplitude. Those below the noise thresholds (grey dots in Fig. 3) were discarded among these data points. In some instances, the above threshold *putative twitches* (coloured dots in Fig. 3) was obtained by combining the same MU and different ultrasound components. In these cases, the combination leading to the highest *putative twitch* was retained (*MU-matched* components, red circles in Fig. 3). The *MU-matched* components had a higher JSC than the *non-MU-matched* (grey dots) components (0.61 ± 0.12 vs 0.26 ± 0.26; *p* < 0.001) (Fig. 3). Noteworthy, the *MU-matched* components had a mean JSC always greater than 0.38, suggesting good repeatability (Fig. 2). In addition, defining the components as *repeatable* using this empirical threshold of 0.38, each recording had 6.5 ± 3.3 repeatable components.

**Figure 3.**
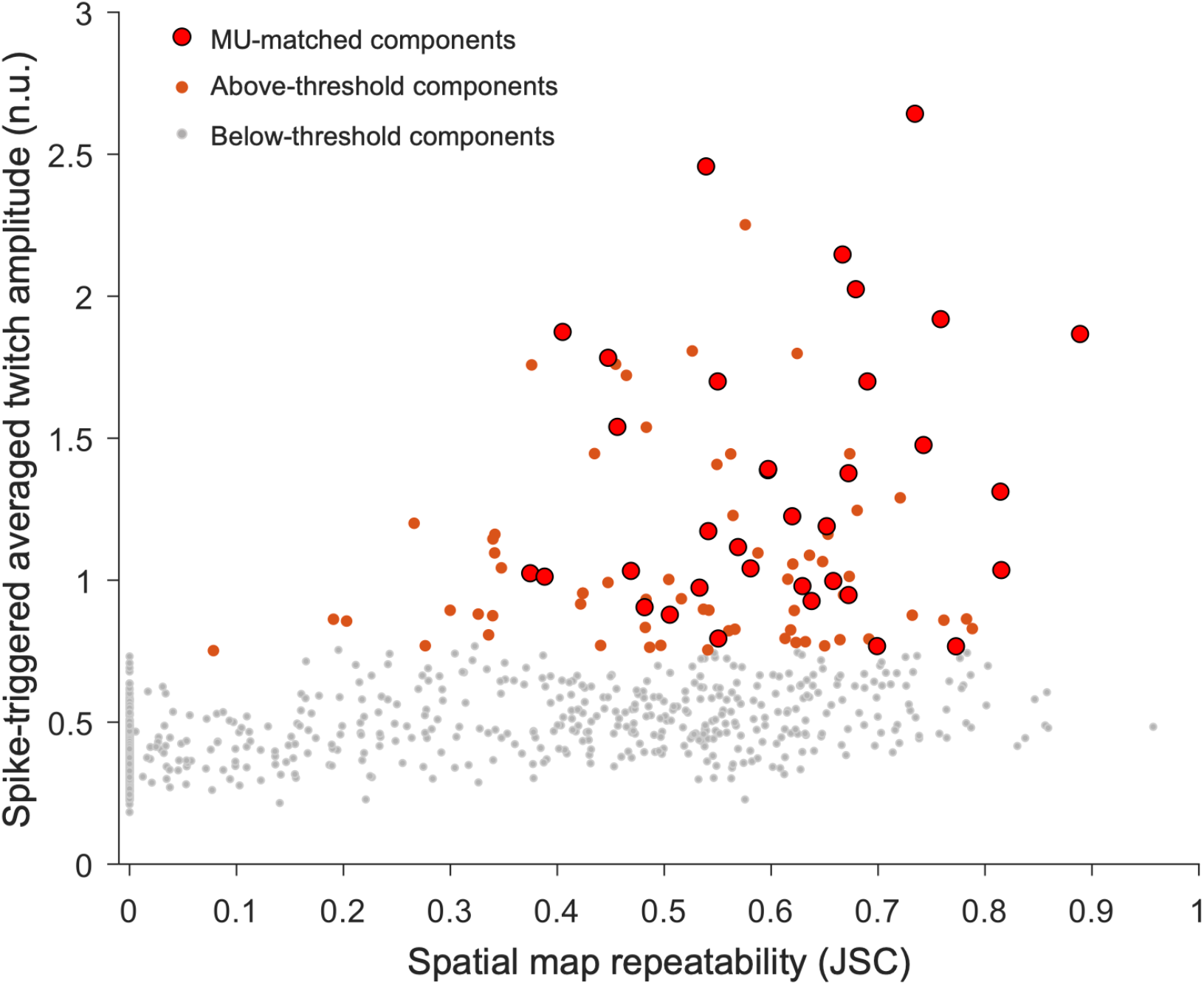
Relationship between Jaccard Similarity Coefficient (JSC) and putative twitches with the highest spike-triggered averaged twitch amplitude. Grey dots are the putative twitches below the noise threshold that were discarded. The red circles correspond to the 35 MU-matched components. All the MU-matched components have JSC over 0.38 (i.e., repeatable). Orange dots refer to multiple components associated with the same MU (e.g., twisting/split territory, duplicate components, etc., see Fig. 5).

Fig. 4 shows three representative examples illustrating the spatial agreement between MUAP distributions and spatial maps of the *MU-matched* components together with the corresponding velocity twitches obtained with spike trigger averaging over all the MU firings of all epochs. *MU-matched* components were spatially (medio-laterally) adjacent to the MUAP distribution (Table 1), as demonstrated by the mediolateral distance between the centroid of the MUAP distributions and the centroid of the spatial maps (5.35 ± 5.17 mm, N = 35 MU). The centroids of the mean spatial maps were distributed across the whole field of view with depths between 2.90 mm and 14.01 mm (Table 1). In addition, the *MU-matched* components had a diameter of 4.03 ± 1.28 mm, similar to previously reported findings of MU territory size using scanning-EMG [32].

**Table 1.** Descriptive statistics about the motor unit-matched repeatable components.

| MU-matched repeatable components | N = 35 |
| --- | --- |
| Jaccard Similarity Coefficient, JSC | 0.61 ± 0.13<br>(0.38; 0.89) |
| Amplitude (n.u) | 1.35 ± 0.49<br>(0.76; 2.64) |
| Centroid-to-centroid (EMG-UUS) (mm) | 5.35 ± 5.17<br>(0.01; 15.83) |
| Depth (mm) | 9.47 ± 2.40<br>(2.90; 14.01) |
| Diameter (mm) | 4.03 ± 1.28<br>(1.45; 7.25) |
| Area (mm²) | 14.06 ± 8.71<br>(1.66; 41.30) |
| Mean ± SD (min; max), MU = motor unit, EMG = electromyography, UUS = ultrafast ultrasound, n.u. = normalised units. |  |

**Figure 4.**
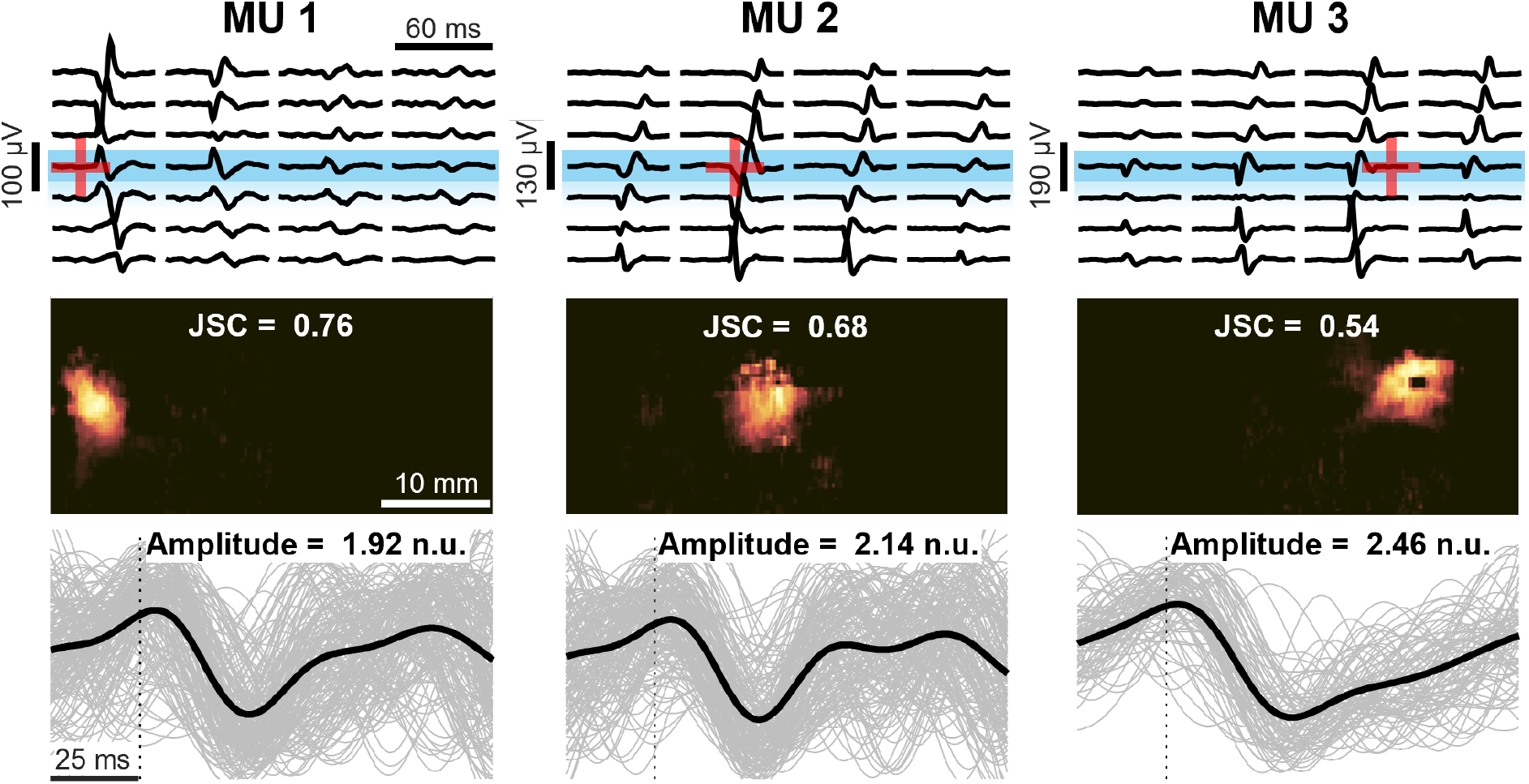
Three representative matches between repeatable components and the motor units (MUs). The upper panels show the MU action potentials and the centroid of the EMG distribution (red ‘+’). In this representation, only the four columns of the EMG grid superimposed on the ultrasound probe (blue rectangle) are shown. The middle panels show the mean spatial map of the repeatable component and the corresponding mean JSC. Finally, lower panels depict the spike-triggered averaged velocity twitch (black line) based on the triggered signals from all seven epochs (grey lines) and the corresponding peak-to-peak amplitude. The vertical dotted lines corresponded to the firing instants of the MUs identified from HDsEMG decomposition and used for the triggering.

### Number of matched components with MUs from HDsEMG: intra and full sequence approach

The intra-sequence analysis led to 35 *MU-matched* components, i.e., 35.4% of the MUs identified by HDsEMG (Table S1, Supplementary material). By decomposing the full eight-second UUS, we found 36 matches, i.e., 36.4% of the MUs identified by HDsEMG. We found no difference in the number of matched MUs across all recordings concerning the two approaches (*p* = 0.9844).

## Discussion

This study investigated whether the spatial repeatability of components extracted from UUS sequences can be used as a criterion to separate muscle tissue displacements associated with single MU activation from noise during stable low-force isometric contractions. First, we decomposed displacement velocity sequences from consecutive two-second epochs of eight-second UUS recordings. Then, we quantified the repeatability of the components’ spatial map across epochs and examined whether there was an association between the repeatability level and the degree of matching with reference MUs identified through HDsEMG decomposition. Finally, we investigated whether this intra-sequence approach using short epochs affects the number of matched MUs by comparing it with the decomposition of the recordings’ full length (eight seconds). We obtained three main findings: 1) all the MU-matched components had a JSC larger than 0.38 and accounted for about one-third of the HDsEMG-detected MU, (2) The components with JSC > 0.38 represented approximately 14% of the 125 initial components from each recording, and (3) the number of MU-component matches did not differ between the intra- and full-sequence approaches.

About 14% of the spatiotemporal components identified applying stICA to UUS sequences were matched with MUs decomposed independently from HDsEMG. A common characteristic of all the *MU-matched* components was the high JSC (Fig. 3) of their spatial maps. This evidence suggests that spatial repeatability across a short epoch is a relevant feature useful to identify putative MUs and implement data reduction methods on the initial set of ultrasound components. This result confirms the initial hypothesis, i.e., since the location of the MU fibres is an invariant feature of the MU during stable isometric contractions, *repeatable* spatial maps are more likely to be associated with actual MUs. Whether this hypothesis applies to conditions other than isometric or constant force contractions likely depends on how MU territory is represented in the ultrasound scanning plane and how this representation changes during a contraction. For instance, muscle shape changes occurring during dynamic contractions may lead to a shift or a shape change of the area where MU fibres’ activation induces movement within the muscle cross-section, i.e., within the ultrasound scanning plane. This would clearly undermine the assumption of MU territory spatial invariance, which is the basis for our hypothesis. Although to a lesser extent, similar variations in MU territory representation can also occur during isometric contractions, for instance, during force-varying contractions, fatiguing contractions or any condition inducing a progressive MU recruitment or de-recruitment. Further studies are required to quantify the effects of these factors on UUS decomposition.

About one-third of MUs decomposed from HDsEMG matched with repeatable ultrasound components. This is similar to the number of successful identifications found in previous studies. It has been previously associated with differences in detection volume and characteristics of two detection systems (EMG and ultrasound) [4,6,33]. In addition to the characteristics of the two measuring techniques, it is worth noting that the measured system is expected to be non-linear due to the heterogenic composition of linear and non-linear elastic constituents. Already at 5-10% MVC, many MUs are active and may suppress or distort the triggered twitch amplitude. Another aspect to consider is that, in this study, we found more repeatable ultrasound components for each recording (6.5 ± 3.3) than HDsEMG MUs (4.9 ± 1.8). Although ultrasound provides a larger field of view and higher spatial resolution than HDsEMG, it remains unclear whether these unmatched repeatable components are MUs and whether they identify different MUs in the whole active MU population. In the present study, the number of successful identifications may have been biased by one subject for which our matching criteria led to no matched MUs. This case was most likely due to the poor quality of the displacement velocity images. The exclusion of this subject would have increased the percentage of MU-matches from 35.4% to 42.7% for the intra-sequence repeatability approach and from 36.4% to 43.9% for the original decomposition over the full sequence (Table S1, Supplementary Material).

Decomposing displacement velocity images into components using stICA over partially overlapping windows likely resulted in component duplicates. Fig. 5a shows two examples of duplicates in which three different components decomposed in three consecutive ROIs showed an amplitude of the twitches (related to the same MU firings) over the noise threshold. In this case, the component providing the highest twitch amplitude was selected and regarded as the MU-matched component. Moreover, it is worth noting that the stICA approach we used assumes spatial independence to decompose the dataset [30,34]. For this reason, it may split MU territories into separate components if the MU activation results in complex movements, e.g. due to the interaction between active and passive tissue [35,36] or tissue rotation due to so-called MU twisting [1,7]. In this regard, Fig. 5b shows two examples of MU twisting of two identified MUs. Two components (matched with the same MU) are spatially separated in two regions of activation (blue and green spots in Fig. 5b) close to each other with inverted twitch shapes (blue and green twitches in Fig. 5b). The shape of the twitch is related to the direction of the movement. In Fig. 5b, the green twitches are negative (i.e., towards the probe/up), while the blue ones are positive (i.e., away from the probe/down). All these examples of duplicate components are now separated and contribute to the above-threshold components in Fig. 3 (small orange points). In future studies, components belonging to the same MU may be merged considering the spatial overlay or a correlation approach based on, e.g., the temporal signals.

**Figure 5.**
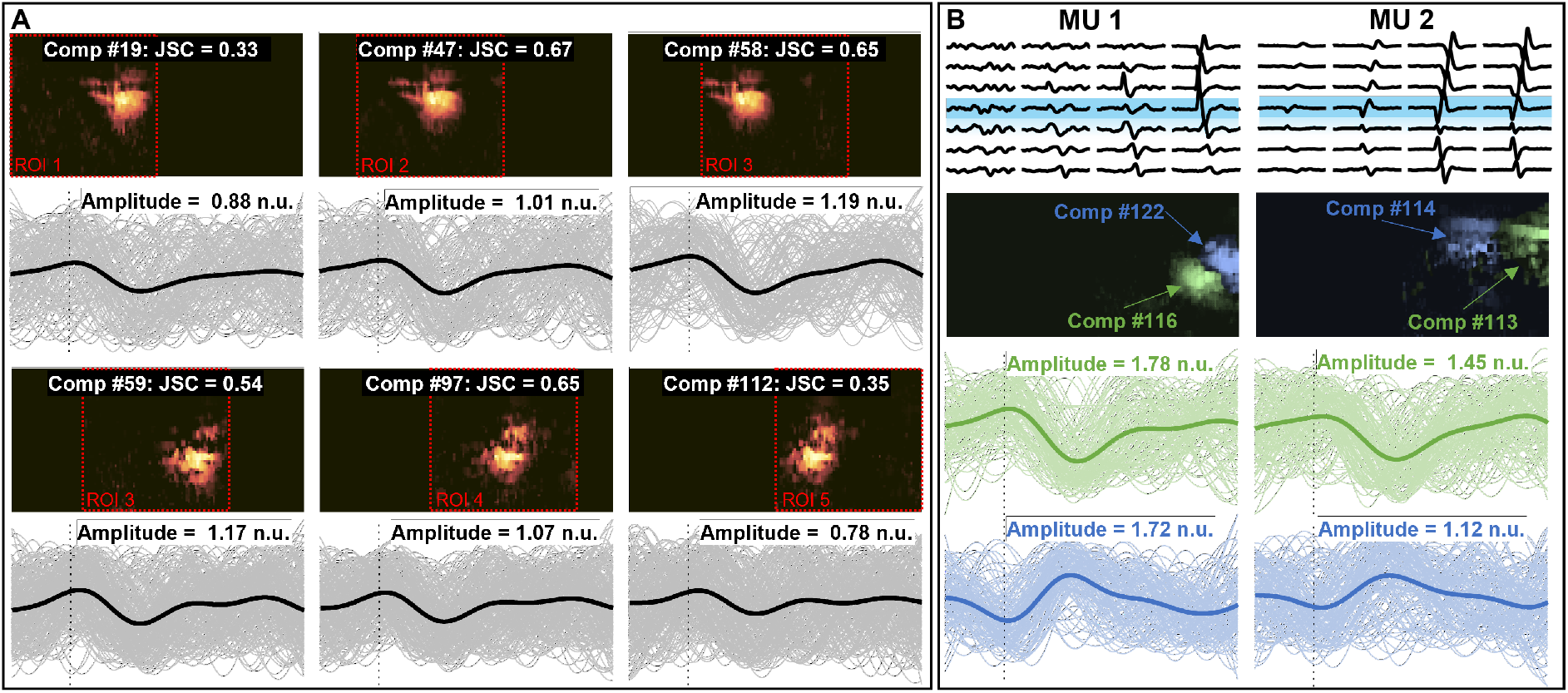
Examples of multiple components associated with the same MU. **A**. Two examples of three different components (belonging to different ROIs) with a similar spatial map (active region) matched with the same MUs. In this case, the three components were merged into the same repeatable component. **B**. Two examples of possible twisting MUs. The MUs were matched with two components showing active regions close to each other and the average twitches showing opposite profiles. Green twitches are negative (movements towards the probe/skin), and blue twitches, on the contrary, are positive (movements away from the probe/skin).

Although finding repeatable components requires eight seconds with the intra-sequence approach herein proposed, the results of this study confirm previous studies that the UUS decomposition method can identify possible MU activity in recordings as short as two seconds [4]. Identifying MUs from a short sequence is an advantage over other methods, such as spike-triggered averaging [9], which requires longer recordings due to other simultaneously active MUs and the motion of non-muscular structures hiding large parts of the movement caused by the target MU. Therefore, the blind source separation approach provides advantages compared to the spike-trigger averaging approach, such as lower memory and storage requirements and a potential to be used for, e.g., real-time imaging [37] and dynamic contractions applications. For these applications, future studies must consider the lower bound in terms of the recording duration to identify MUs and improve the classification of components into MUs or non-MUs using robust features or training a classifier. For example, the Gaussian-like 2D distribution of velocities reported in this work for the most repeatable components and similar to what has been found in previous studies [1,4,6,7], may be a feature for the classification of a component as a MU. Thus, having a classifier for MU/non-MU-associated components enables the UUS approach to be stand-alone from HDsEMG.

In conclusion, this study investigated the association of intra-sequence repeatable components with individual MU activity. We found that 1) spatial repeatability can be used as a data reduction to select putative MU activity during stable isometric contractions, and 2) the UUS decomposition method can identify possible MU activity in two-second recordings equally well as in eight-second recordings. These findings provide a foundation for developing stand-alone methods to identify MU in ultrafast ultrasound and represent a step towards real-time imaging of active MU territories.

## Acknowledgements

RR is supported by the Swedish Research Council for Sport Science (grant number: D2023-0003). MC is supported by the project “Trajector-AGE” (grant number: 2020477RW5PRIN), funded by the Italian Ministry of Universities and Research. CG is supported by the Swedish Research Council (grant number: 2022-04747).

