## Supplementary material for "Spatially repeatable components from ultrafast ultrasound are associated with motor unit activity in human isometric contractions"

**Table S1.** Descriptive statistics about the recordings, decomposed motor units (MUs) and repeatable components ('US-RepMap', components with JSC over 0.38). Duplicates and splitting components were counted as one component in the 'Without duplicates' column.

| Trials | Subject | MU | US-EMG match | % of US-EMG match | 8s US-EMG match | 8s % of US-EMG match | US-RepMap | % of US-RepMap | Without duplicates |
|---|---|---|---|---|---|---|---|---|---|
| **1** | **S1** | 10 | 3 | 30.0% | 5 | 50.0% | 21 | 16.8% | 9 |
| **2** | **S1** | 3 | 2 | 66.7% | 2 | 66.7% | 15 | 12.0% | 6 |
| **3** | **S1** | 6 | 0 | 0.0% | 1 | 16.7% | 13 | 10.4% | 4 |
| **4** | **S1** | 3 | 2 | 66.7% | 2 | 66.7% | 18 | 14.4% | 4 |
| **5** | **S2** | 5 | 4 | 80.0% | 4 | 80.0% | 21 | 16.8% | 7 |
| **6** | **S2** | 7 | 0 | 0.0% | 1 | 14.3% | 22 | 17.6% | 7 |
| **7** | **S2** | 4 | 1 | 25.0% | 2 | 50.0% | 19 | 15.2% | 9 |
| **8** | **S2** | 4 | 1 | 25.0% | 1 | 25.0% | 19 | 15.2% | 8 |
| **9** | **S2** | 4 | 2 | 50.0% | 2 | 50.0% | 15 | 12.0% | 7 |
| **10** | **S3** | 5 | 2 | 40.0% | 4 | 80.0% | 8 | 6.4% | 2 |
| **11** | **S3** | 3 | 3 | 100.0% | 1 | 33.3% | 20 | 16.0% | 6 |
| **12** | **S4** | 8 | 6 | 75.0% | 4 | 50.0% | 28 | 22.4% | 9 |
| **13** | **S4** | 5 | 3 | 60.0% | 3 | 60.0% | 26 | 20.8% | 14 |
| **14** | **S4** | 6 | 2 | 33.3% | 2 | 33.3% | 24 | 19.2% | 11 |
| **15** | **S4** | 4 | 1 | 25.0% | 0 | 0.0% | 13 | 10.4% | 4 |
| **16** | **S4** | 5 | 3 | 60.0% | 2 | 40.0% | 23 | 18.4% | 10 |
| **17** | **S5** | 4 | 0 | 0.0% | 0 | 0.0% | 22 | 17.6% | 7 |
| **18** | **S5** | 5 | 0 | 0.0% | 0 | 0.0% | 8 | 6.4% | 2 |
| **19** | **S5** | 5 | 0 | 0.0% | 0 | 0.0% | 11 | 8.8% | 3 |
| **20** | **S5** | 3 | 0 | 0.0% | 0 | 0.0% | 5 | 4.0% | 1 |
| | **Total** | **99** | **35** | **35.4%** | **36** | **36.4%** | **351** | | **130** |
| | **Average** | **4.95** | **1.75** | **36.8%** | **1.8** | **35.8%** | **17.55** | **14.0%** | **6.5** |
| | **SD** | **1.8** | **1.6** | **31.6%** | **1.5** | **27.7%** | **6.3** | **5.1%** | **3.3** |
| | Without S5 | **82** | | **42.7%** | | **43.9%** | | | |

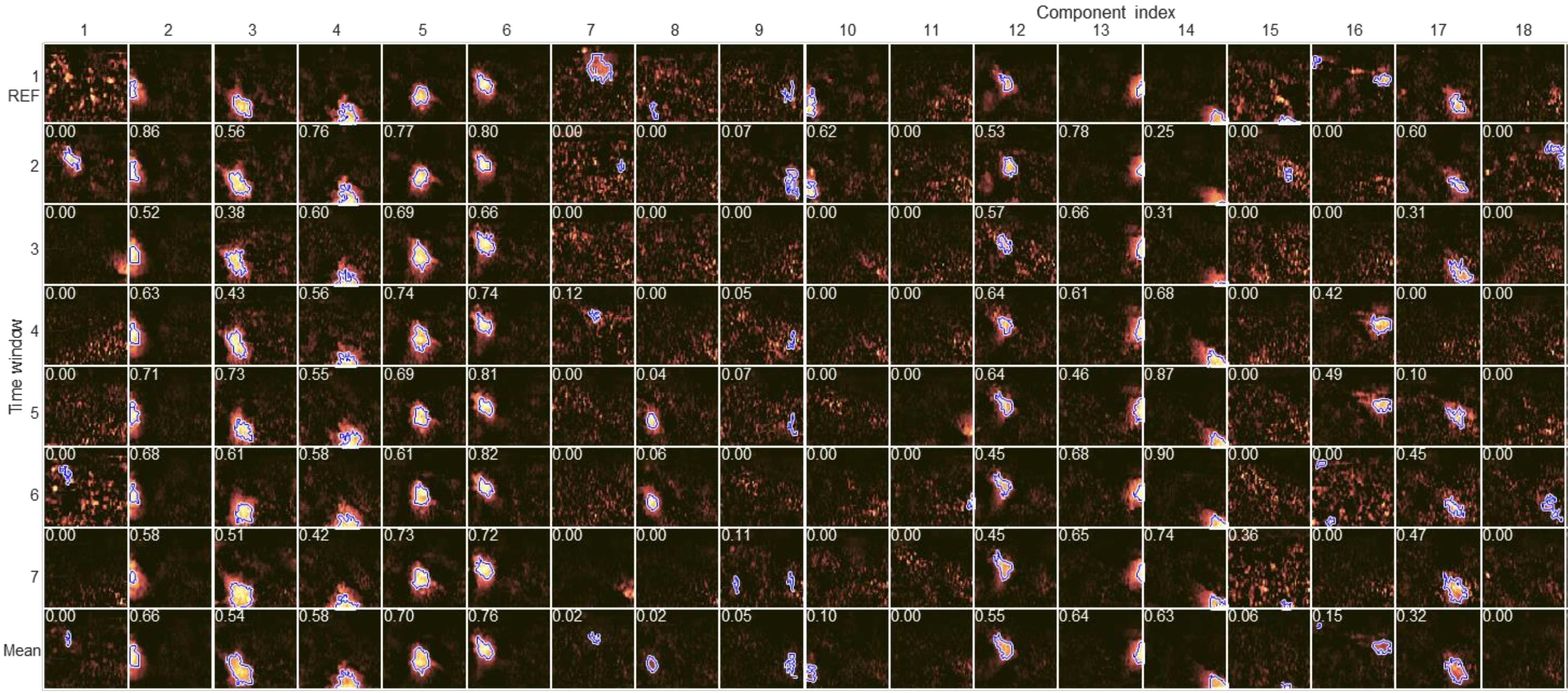


**Figure S1.** Example of components (columns) over the seven epochs (rows) with the corresponding Jaccard Similarity Coefficient (JSC) calculated with respect to the reference first epoch. The last row shows the average of the averaged spatial map. The blue lines represent the territories identified in the images with the k-means. This figure shows the large variability of JSC values.
